# Global and local connectivity differences converge with gene expression in a neurodevelopmental disorder of known genetic origin

**DOI:** 10.1101/057687

**Authors:** Joe Bathelt, Jessica Barnes, F Lucy Raymond, Kate Baker, Duncan Astle

**Affiliations:** MRC Cognition & Brain Sciences Unit, Cambridge, UK; Department of Medical Genetics, Cambridge Institute for Medical Research, University of Cambridge, Cambridge, United Kingdom

## Abstract

Knowledge of genetic cause in neurodevelopmental disorders can highlight molecular and cellular processes critical for typical development. Furthermore, the relative homogeneity of neurodevelopmental disorders of known genetic origin allows the researcher to establish the subsequent neurobiological processes that mediate cognitive and behavioural outcomes. The current study investigated white matter structural connectivity in a group of individuals with intellectual disability due to mutations in *ZDHHC9*. In addition to shared cause of cognitive impairment, these individuals have a shared cognitive profile, involving oro-motor control difficulties and expressive language impairment. Analysis of structural network properties using graph theory measures showed global reductions in mean clustering coefficient and efficiency in the *ZDHHC9* group, with maximal differences in frontal and parietal areas. Regional variation in clustering coefficient and local efficiency across cortical regions in cases and controls were significantly associated with known pattern of expression of *ZDHHC9* in the normal adult human brain. The results demonstrate that a mutation in a single gene impacts upon white matter organisation across the whole-brain, but also shows regionally specific effects, according to variation in gene expression. Furthermore, these regionally specific patterns may link to specific developmental mechanisms, and correspond to specific cognitive deficits.

## 2 Introduction

Many cognitive and psychiatric disorders are highly heritable (Lee 2013, Sullivan 2012, Haworth 2009). In some cases, genetic risk factors have been identified, but understanding the neural mechanisms linking altered gene transcripts to cognitive or behavioural outcomes remains challenging. One reason for this is the heterogenous nature of the vast majority of these disorders, which presents a major challenge to establishing the neural endophenotypes that mediate any gene-cognition relationships; any group defined on the basis of a cognitive impairment or behavioural difficulty will likely contain individuals with different genetic and neural causes, making it difficult to identify mechanisms at the group level. One promising approach has been to study brain organisation in groups of individuals that have rare but clearly defined genetic causes of those impairments (Meyer-Lindenberg 2009, Griffa 2013). These groups, whilst necessarily small in size, have an homogenous aetiology. Studying these groups can therefore provide a powerful means for identifying the neurobiological pathways that potentially mediate cognitive and behavioural phenotypes in the wider population. For instance, the study of a rare familial speech disorder (KE family, *F0XP2* mutation) highlighted the importance of striatal networks for emergent higher-order language skills (Watkins 2011, Liégeois 2011).

However, studies of brain differences have mainly focussed on focal differences in brain areas or white matter tracts that show the most pronounced group differences. This is true of both genetically defined group comparisons and case-control designs more generally. However, genetic differences are likely to have wide-ranging effects on the organisation of neural ensembles across many areas. To explore this fully requires a more advanced network science approach, cable of establishing how organisational principles differ across groups of individuals (Petersen 2015, Meyer-Lindenberg 2009). We take this approach here.

In a network analysis brain regions are described as a nodes and their connections as edges. Nodes typically correspond to regions of interest (Dell’Acqua 2012, Fornito 2015). Edges can represent the strength of white matter connectivity based on diffusion-weighted imaging (Qi 2015). Graph theory provides a mathematical framework for the analysis of the resulting network (Bullmore 2009, Rubinov 2010), which describes organisational principles related to ease of information exchange and wiring cost. Structural network approaches have been used to study typical and atypical brains across the lifespan (Griffa 2013, Hagmann 2012, Di Martino 2014, Deco 2014). These studies have revealed an organisational principle of high local connectivity with some additional long-range connections (Bassett 2006). This organisation provides an optimal trade-off between ease of information transfer and minimization of wiring costs (Fornito 2015).

White matter has been shown to be highly heritable indicating a high level of genetic influence on white matter structure (Lee 2015, Kochunov 2015). A few studies have employed a network analysis approach to investigate if this also extends to structural network organisation (Ottet 2013, Meoded 2014, Hong 2014, Leow 2014, Bruno 2016). These studies focussed on common variants of trophic factor genes (Meoded 2014), genes involved the regulation of synaptic weights (Ottet 2013, Meoded 2014, Hong 2014), and mutations associated with specific phenotypes (Leow 2014, Bruno 2016). Genetic differences were associated with differences in structural brain network organisation, with specific effects for each genetic factor. This suggests that studying differences in brain organisation may offer important insight into understanding the effects of genetic variation.

In the present study we take a network analysis approach to studying brain organisation in a genetically defined neurodevelopmental disorder. Mutations in *ZDHHC9* are a recurrent cause of X-linked Intellectual Disability (XLID) (Raymond 2007). The *ZDHHC9* gene codes for a palmitoylation enzyme, involved in post-translational modification of specific target substrates. Palmitoylation plays an important role in the recruitment of receptors and ion channels at the synapse (Topinka 1998, Young 2013, El-Husseini 2000). A systematic assessment of clinical history and cognitive deficits across multiple XLID-associated genes led to the observation that *ZDHHC9* mutations are associated with homogeneous neurological and cognitive features, including disproportionate attention problems, language impairment, and deficits in oromotor control in the context of mild to moderate intellectual disability (Baker 2015). The majority of affected individuals also had a history of epilepsy that resembled Rolandic epilepsy in presentation and spike topography (Baker 2015). Previous neuroimaging work in our group investigated focal differences in brain structure in *ZDHHC9* cases. These studies indicated differences in subcortical volumes (thalamus, putamen, caudate nucleus) and hypoplasia of the corpus callosum (Baker 2015). Reductions in cortical thickness were found that were most pronounced in areas around the temporo-parietal junctions and inferior frontal lobe (Bathelt et al. *under review*). Mutation of *ZDHHC9* was also associated with reductions in white matter structural integrity involving cortical, cortico-subcortical, and interhemispheric connections (Bathelt et al. *under review*).

Given strong evidence for pervasive effects on white matter integrity, we expected the *ZDHHC9* mutation to influence information transfer within the brain network More specifically, we predicted that in addition to any global impact of gene mutation, we ought to observe some regional specificity, according to variability in the expression of that gene across the brain. This regional specificity may correspond to the areas of most marked cognitive impairment resulting from the mutation, and overlap with other genes known to impact upon similar developmental mechanisms. In short, across our analyses we explored how both a mutation to, and regional expression of, *ZDHHC9* are associated with structural brain organisation.

## 3. Methods

### 3.1 Participants

The study compared 7 males with inherited loss of function mutations in the *ZDHHC9* gene (Age in years: mean=29.13, SE=4.86, Range=13.83−41.83) to 7 males individually matched in age +/− 2 years (Age in years: mean=27.23, SE=5.31, Range=10.17-42.5). Comparison subjects had no history of neurological illness or cognitive impairment. Statistical analysis indicated no significant difference in age between the groups (Welch-corrected t-test: t(11.91)=−0.265, p=0.796).

For detailed description of clinical and cognitive characteristics of the *ZDHHC9* group see Baker et al. 2015. In summary, all individuals with a *ZDHHC9* mutation had mild to moderate intellectual disability (full-scale IQ: mean=64.86, SE=2.32, Range=57-73). 5 individuals had a history of epilepsy, with seizure characteristics and EEG features similar to the Rolandic epilepsy spectrum. At the time of MRI acquisition, 1 participant reported seizures within the previous 3 months, and 3 currently received anti-epileptic medication (carbemazapine n=l, carbemazapine and lamotrigine n=1, phenytoin n=1). Vineland scores (Sparrow 2005) indicated stronger receptive language abilities compared to expressive and written language abilities in the *ZDHHC9* group. The Verbal Motor Production Assessment for Children (VMPAC) (Hayden 1999) indicated significant oromotor difficulties in the *ZDHHC9* group, including deficits in oral control, sequencing, voice characteristics, and connected speech. Inhibitory control was also reduced in the *ZDHHC9*group on a visual attention task. These specific features differentiated with *ZDHHC9* group from age and IQ matched controls (Baker 2015).

### 3.2 MRI data acquisition

Magnetic resonance imaging data was acquired at the MRC Cognition and Brain Sciences Unit, Cambridge U.K. All scans were obtained on the Siemens 3 T Tim Trio system (Siemens Healthcare, Erlangen, Germany), using a 32-channel quadrature head coil. The imaging protocol consisted of two sequences: T1-weighted MRI and a diffusion-weighted sequence.

T1-weighted volume scans were acquired using a whole brain coverage 3D Magnetisation Prepared Rapid Acquisition Gradient Echo (MP RAGE) sequence with 1mm isometric image resolution. Echo time was 2.98 ms, and repetition time was 2250 ms.

Diffusion scans were acquired using echo-planar diffusion-weighted images with an isotropic set of 60 non-collinear directions, using a weighting factor of b = 1000 s/mm^−2^, interleaved with 4 T2-weighted (b = 0) volumes. Whole brain coverage was obtained with 60 contiguous axial slices and isometric image resolution of 2mm. Echo time was 90 ms and repetition time was 8400 ms.

### 3.3 Structural connectome analysis

The white-matter connectome reconstruction followed a commonly used procedure of estimating the most probable white matter connections for each individual, and then obtaining measures of fractional anisotropy (FA) between cortical regions after transformation to common space (Horn 2016). Computer code for this workflow and subsequent analyses is available online (https://github.com/joebatheIt/ZDHHC9connectome). The details of the procedure are described in the following paragraphs.

In the current study, MRI scans were converted from the native DICOM to compressed NIfTI-1 format using the dcm2nii tool developed at the McCauseland Centre for Neuroimaging http://www.mccau.slandcenter.sc.edu/mricro/mricron/dcm2nii.html. Subsequently, the images were submitted to an implementation of a non-local means de-noising algorithm (Coupe 2008) in the Diffusion Imaging in Python (DiPy) v0.8.0 package (Garyfallidis 2014) to boost signal to noise ratio. Next, a brain mask of the b0 image was created using the brain extraction tool (BET) of the FMRIB Software Library (FSL) v5.0.8. Motion and eddy current correction were applied to the masked images using FSL routines. The corrected images were re-sliced to 1mm resolution with trilinear interpolation using in-house software based on NiBabel v2.0.0 functions http://nipy.org/nibabel/. A spherical constrained deconvolution (CSD) model was fitted to the 60 gradient direction diffusion-weighted images using a maximum harmonic order of 8. Correct anatomical orientation of CSD glyphs was visually inspected for white matter tracts of known orientation (corpus callosum, cortico-spinal tract).

A whole-brain tractography was generated by seeding streamlines from each voxel within the brain mask using the probabilistic tractography algorithm of MRTrix (Tournier 2012). The desired number of streamlines was set to 150,000. Other settings followed the recommendations for MRTrix: The fibre tracking algorithm was set to a minimum and maximum track length of 10mm and 200mm respectively. The minimum radius of curvature was set to 1 mm and the track size to 0.2mm. The track termination threshold was set to an FA value of 0.1.

Subsequently, a 12 degree-of-freedom affine transform between each participant’s skull-stripped FA image and the FMRIB58 template in MNI152 was calculated with the correlation ratio as the cost function using FSL FAST. This transform was applied to the streamlines in each participant’s anatomical space to move them into MNI common space (Horn 2016) using a TrackVis v0.6.01 algorithm (Wang et al., 2007).

For structural connectome analysis, regions of interests (ROIs) were based on the Desikan-Killiany parcellation of the MNI template (Klein & Tourville 2012) with 34 ROIs per hemisphere. The ROIs filled the space between the cortical grey and white matter so that streamlines would terminate at the edges of the ROI. For each pairwise combination of ROIs, the number of streamlines intersecting both ROIs was estimated and transformed to a density map. This density map was binarized and multiplied with the FA map to obtain the FA value corresponding to the connection between the ROIs. This procedure was implemented in-house based on DiPy v0.8.0 functions (Garyfallidis 2014). Only cortical ROIs were considered in the current analysis, because Allen Brain atlas data could only be mapped for these ROIs (French & Paus, 2015). Visualizations of the structural connectome were generated using the BrainNet Viewer toolbox (Xia et al.,2013).

### 3.4 Graph theory

Structural networks were analysed using graph theory metrics to compare network characteristics between the groups and relate node properties to the expression profile of *ZDHHC9*. Graph metrics were calculated in the python implementation of the Brain Connectivity Toolbox (https://sites.google.com/site/bctnet/). Weighted undirected networks were used for all analyses. The weight represented the FA value in the structural connectome. A detailed description of the graph theory metrics used in the analysis can be found elsewhere (Bullmore & Sporns, 2009, Rubinov & Sporns, 2010). In brief, for characterisation of network nodes, node degree, node strength, and clustering coefficient were used. The node degree is the number of edges connected to a node. The node strength is the sum of the weight of edges connected to a node. The clustering coefficient is the number of connections of a node to its neighbours relative to all possible connections. For a characterisation of the global network, global efficiency was calculated. Global efficiency describes the ease of information transfer within a network based on the number of nodes that have to be traversed to reach any node from any other node in the network.

### 3.5 Allen Brain Atlas data

Gene expression data were obtained from the Allen Brain Atlas Human Brain public database (http://human.brain-map.org). Gene expression data were based on microarray analysis of post-mortem tissue samples from 6 human donors between 18 and 68 years with no known history of neuropsychiatric or neurological conditions (see online documentation). MRIs and transformations from individual donors MR space to MNI coordinates were also obtained from the Allen Brain Atlas website. For the current investigation, expression values were averaged across donors and mapped onto areas of the Desikan-Killiany parcellation of the MNI brain as described by French and colleagues (French & Paus, 2015). The current investigation focussed on the expression of *ZDHHC9*. In order to investigate the specificity of the link of *ZDHHC9* expression and structural connectome organisation, we compared *ZDHHC9* to a number of other genes: First, *GAPDH* was added as a control gene that is not associated with any known neurological or cognitive phenotype (Nicholls et al., 2012). We then assessed genes that are associated with a similar mutation phenotype. For overlap with language deficits, *F0XP2* was included (Khadem et al., 2005). *FMR1* was selected as an X-linked intellectual disability gene (Bourgeois et al., 2009). *GRIN2A* was included for the association with Rolandic Epilepsy (McTague et al., 2015).

## 4 Results

### 4.1 Structural connectome analysis

Brain networks were obtained from structural connectome descriptions of diffusion-weighted MR data. Adjacency matrices represented the FA value associated with streamlines between pairs of regions in the Desikan-Killiany atlas. A visual representations of the group-average structural connectome is shown in Figure 1. Principles of organisation within the structural networks were analysed using graph theory. Statistical comparison indicated significant brain-wide differences in mean node strength, mean clustering coefficient, and global efficiency with reductions in all measures in the *ZDHHC9* group (see Table 1). There was no statistically significant difference in mean node degree between groups.

**Figure 1:**
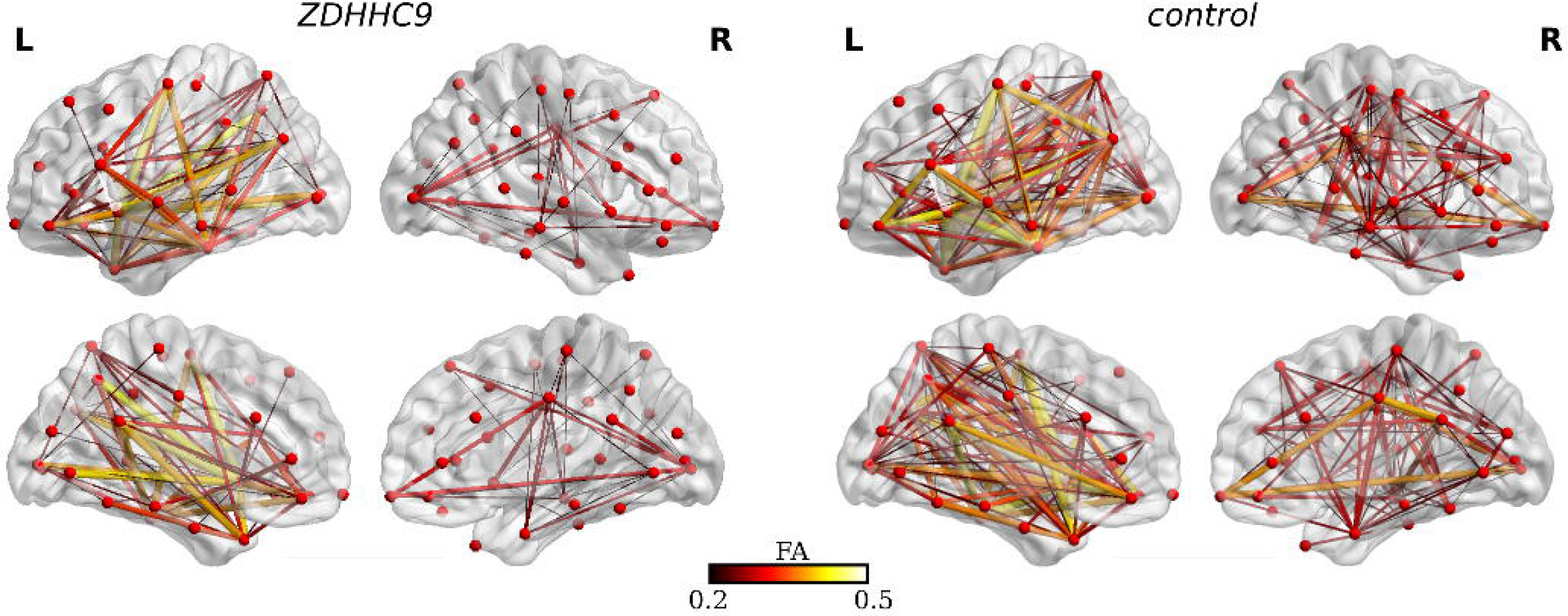
White matter network structure in the ZDHHC9 and control group. The colour values represent the edge weight, i.e. the fractional anisotropy (FA) value associated with a connection between any two ROIs. The adjacency matrix was thresholded at 0.2 for visualization.

Subsequently, local differences in graph theory metrics within cortical regions of the Desikan-Killiany atlas were compared between the *ZDHHC9* and control group. The results indicated significant reductions in the *ZDHHC9* group for the clustering coefficient of the left inferior frontal, middle frontal, and orbito-frontal cortex; precentral, and supramarginal gyrus; and posterior, and isthmus cingulate cortex. For the right hemisphere, reductions in cluster coefficient in the *ZDHHC9* group were found in the supramarginal gyrus, superior temporal gyrus, insula, inferior frontal cortex, lingual cortex, and cuneus. The clustering coefficient of a node is the number of connections to the node’s neighbours relative to the possible number of connections (Rubinov & Sporns, 2011). A reduction in the clustering coefficient of nodes in the *ZDHHC9* group therefore indicates a reduction in the integration of these nodes. A comparison of the topographical pattern of differences in local efficiency indicated widespread reduction in the *ZDHHC9* group expected due to the relationship in the algorithms for calculating clustering coefficient and local efficiency (Rubinov & Sporns, 2011). Regional analysis of node strength and node degree did not show any similar pattern.

**Figure 2:**
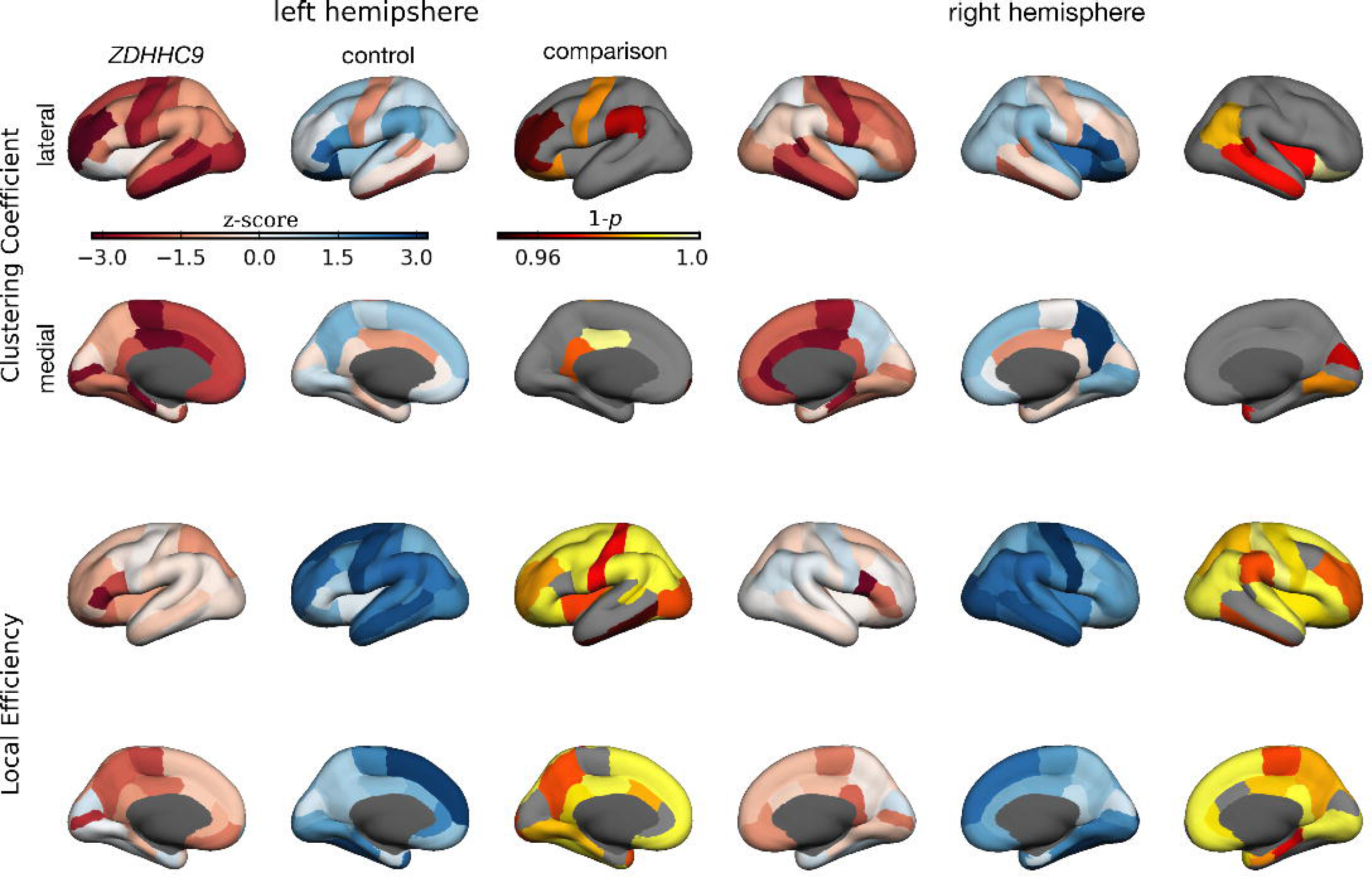
Overview of regional differences in clustering coefficient and local efficiency between the ZDHHC9 and control group. Red-blue maps show the distribution of graph metrics z-scored across hemispheres and participant groups. Yellow-red maps show the results of t-test comparisons of graph metrics corrected for multiple comparison using Bonferroni correction.

**Table 1.**
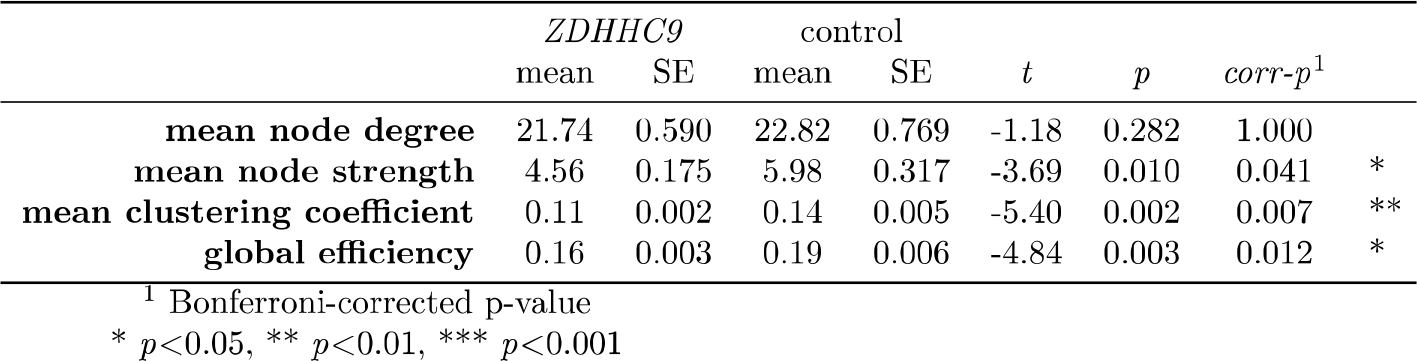
Comparison of global graph theory metrics between the ZDHHC9 and control group. Statistical comparison indicated significantly lower mean node strength, mean clustering coefficient, and global efficiency in the *ZDHHC9* group.

### 4.2 *ZDHHC9* expression and structural connectome properties

We then investigated the relationship between measures of structural connectivity and *ZDHHC9* expression. First, gene expression information was obtained from the Allen Brain Institute Human Brain database and mapped onto the Desikan-Killiany parcellation of the MNI brain (French & Paus 2015). *ZDHHC9* expression was found to be higher in the left compared to the right hemisphere (see Figure 2). Local maxima were found in the left post-central gyrus, inferior frontal cortex, anterior cingulate cortex, inferior parietal lobule, and right lingual gyrus. Low expression was observed in the right posterior and isthmus cingulate cortex, and left superior temporal gyrus.

Next, we investigated whether the variation of *ZDHHC9* expression over cortical regions in the healthy adult brain explained the regional variation of graph theoretical measures in the structural connectome derived from case control analysis of patients with *ZDHHC9* mutations. The Shapiro-Wilk test indicated significant departures from the normality assumption for all variables (*ZDHH9* expression: W=0.98, p=0.071; Node strength: W=0.93, p=0.001; Clustering coefficient: W=0.96, p=0.001; Betweenness centrality: W=0.89, p=0.001; Local Efficiency: W=0.95, p=0.001). Therefore, all variables were transformed to z-scores for further analysis; these z-scores were normally distributed. We firstly checked the correlations between graph measures, which were not sufficiently high to introduce multiple co-linearity problems (all r<0.71). Separate multiple linear regression models were fitted for each graph measure using *ZDHHC9* expression and participant group as predictors. A significantly better fit than the null model was indicated for all regression models, apart from model for betweenness centrality (Node strength: F(2,129)=9.56, p=0.001, R^2^=0.129; Clustering coefficient: F(2,129)=50.91, p=0.001, R^2^=0.441; Betweenness centrality: F(2,129)=0.52, p=0.597, R^2^=0.008; Local efficiency: F(F(2,129)=225.8, p=0.001, R^2^=0.778). A significant effect of participant group was found for node strength, clustering coefficient, and local efficiency (Node strength: t=−4.36, p=0.001; Clustering coefficient: t=-9.73, p=0.001; Local efficiency: t=−21.12, p=0.001). Further, there was a significant effect of *ZDHHC9* expression on clustering coefficient (t=2.67, p=0.009). The findings indicated that cortical areas with higher expression of *ZDHHC9* were linked by white matter with higher FA. This relationship was similar in both groups, but was associated with overall lower graph measure scores in the *ZDHHC9* group.

Lastly, we investigated the specificity of the association between *ZDHHC9* expression and structural connectome graph measures by repeating the analysis with a control brain-expressed gene and with genes with a similar mutation-associated phenotype. As in the main analysis, regression analysis was applied with the graph measure as the outcome and gene expression and participant group as predictors. *F0XP2* showed a significant negative association with clustering coefficient in both groups (β=−0.251, t=−3.93, p<0.001). There were no other significant effects for any graph measure and gene expression combination
.

**Figure 3:**
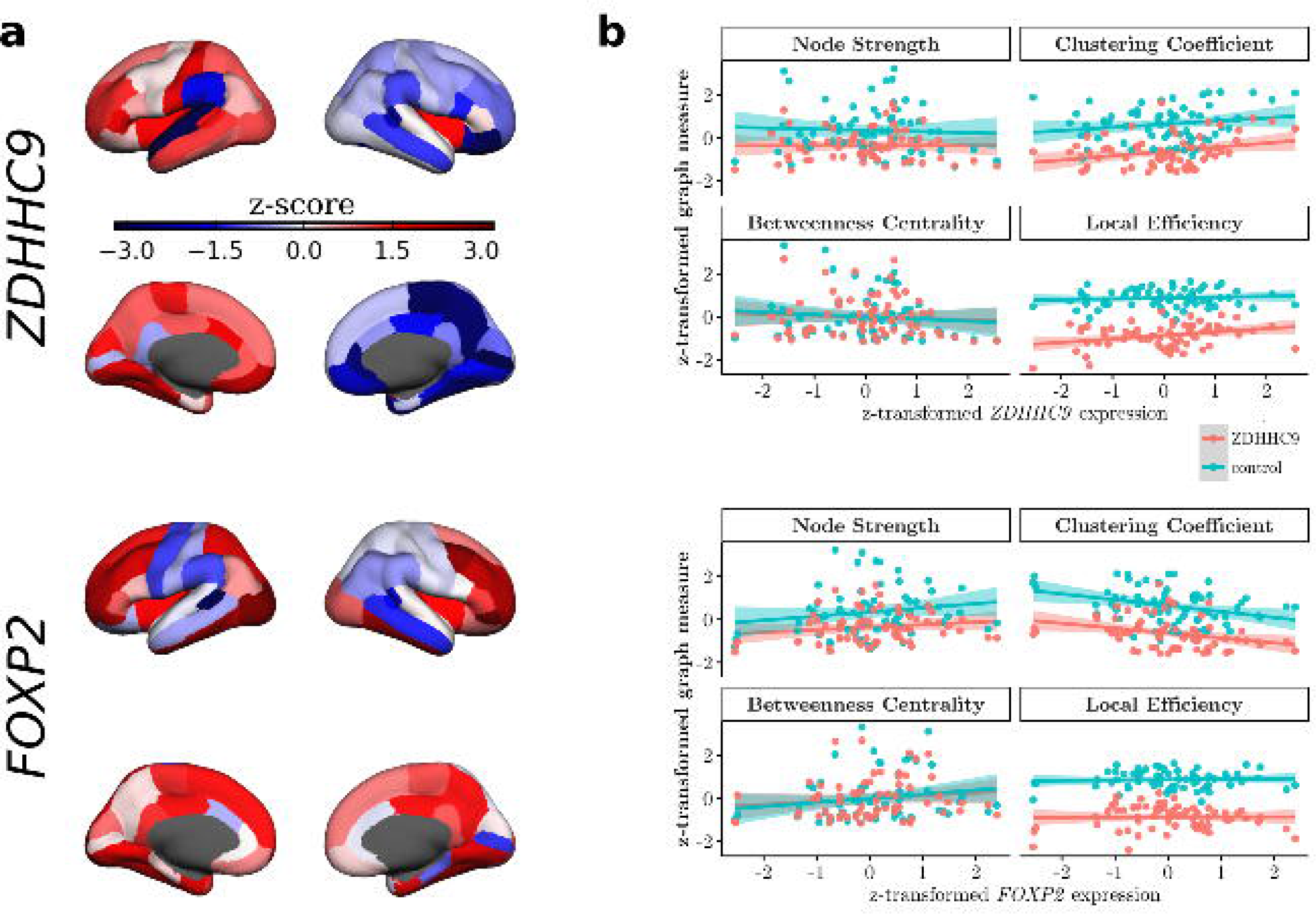
**a**: Expression of *ZDHHC9* (*top*) and *F0XP2* (*bottom*) in regions of the cortex in the normal adult human brain (Allen Brain Atlas), **b**: Relationship between regional variation in *ZDHHC9* (*top*) and *F0XP2* (*bottom*) expression and graph theory measures in the *ZDHHC9* case (*red*) and control (*blue*) groups. Regression analysis indicated a significant effect of participant group for node strength, clustering coefficient, and local efficiency. Local *ZDHHC9* expression was found to be an additional significant predictor for the clustering coefficient and local efficiency. *F0XP2* emerged as a significant negative predictor for regional variation in clustering coefficient.

## 5 Discussion

We investigated the influence of mutations in the palmitoylation-enzyme-gene *ZDHHC9* on differences in white-matter-network organisation. To this end, a connectome was constructed to describe the fractional anisotropy (FA) value associated with all connections between cortical regions in the Desikan-Killiany cortical parcellation. Comparison of global graph theory measures indicated significantly lower global efficiency accompanied by reductions in mean clustering coefficient and mean node strength. Topological analysis indicated that differences in clustering coefficient were localised to frontal, temporal and parietal areas, whereas differences in local efficiency were more pervasive.

Further, we investigated the relationship between *ZDHHC9* expression and structural connectivity of cortical regions. Analysis of the relationship between regional variation in *ZDHHC9* expression and structural network metrics in both groups indicated a significant effect of clustering coefficient with higher expression being associated with a higher clustering coefficient. In contrast, regional variation in *F0XP2* expression showed a negative association with clustering coefficient, while other genes with overlapping phenotypes did not show an association with regional differences in graph metrics.

### *ZDHHC9* mutations result in global changes in brain organisation

In the current investigation we observed global reductions in edge weight in the *ZDHHC9* case group. Edge weight represented the fractional anisotropy (FA) of white matter pathways in the networks. FA is commonly interpreted as a measure of white matter integrity. Reduction in FA may be caused by differences in axon diameter, packing density, or myelination (Alexander 2011). The global reduction in FA mirrors previous results with tract-based spatial statistics analysis (Bathelt et al., *under review*) that indicated widespread differences in white matter integrity in the *ZDHHC9* group.

Graph theoretical analysis indicated that reduced FA in the *ZDHHC9* group was associated with lower mean clustering coefficient and global efficiency. Lower mean clustering coefficient suggests reduced local connectivity of nodes with their neighbours. Reduced global efficiency is linked to reduced information transfer across the structural brain network in the *ZDHHC9* group.

These observed global differences in structural brain network organisation may arise early in brain development. Global network efficiency has been found to be stable across the human lifespan in typical development. The structural core of the white matter connectome is formed during the second to third trimester of pregnancy (Collin 2013). Any differences in genetic predisposition affecting this processes are likely to impose early constraints with general effects on the later trajectory of neural and cognitive development and may contribute to the mild-moderate intellectual disability observed in this group (Di Martino 2014).

### *ZDHHC9* mutations result in regionally-specific changes in brain organisation

Topographical analysis of clustering coefficient and local efficiency indicated differences in nodes of the frontal, left parietal, and right temporal lobe; these nodes are less integrated with the rest of the network in the *ZDHHC9* group. These regionally specific effects may provide a basis for the profile of disproportionate speech and language deficits observed in these individuals (Baker et al 2015).

Furthermore, the expression of *ZDHHC9* was related to the observed regional variation in graph metrics. Regression analysis indicated that a significant effect of *ZDHHC9* expression on the local efficiency and clustering coefficient of nodes. Higher expression of *ZDHHC9* was related to a higher clustering coefficient and higher local efficiency in the *ZDHHC9* mutated group and typical control. Expression of *ZDHHC9* was highest in left parietal and frontal regions that also showed the largest reduction in regional comparison of clustering coefficient and local efficiency in the *ZDHHC9* group. In contrast, regional variations in clustering coefficient was negatively related to the expression of *F0XP2*, a transcription factor implicated in language function (Khadem, 2005), but showed no relationship to the expression of other genes with overlapping mutation phenotype. There are a number of possible explanations for the observation of parallel (but opposite) associations between connectivity and *ZDHHC9* or *F0XP2* expression. Firstly, the observation could be explained by a similar expression topology for both genes, with no direct mechanistic link. However, data from the Allen Brain Atlas does not support this explanation (*ZDHHC9* and *F0XP2* expression region-by-region correlation −0.0445). Alternatively, the observation could suggest that these genes have convergent influence on cortical architecture and cognitive development via a shared molecular pathway. FOXP2 is a transcriptional repressor, hence FOXP2 might down-regulate *ZDHHC9* expression leading to reduced palmitoylation and disrupted trafficking of relevant substrates. Vernes and colleagues (Vernes et al., 2011) found that *ZDHHC3* is regulated by FOXP2, but there is currently no positive evidence that *ZDHHC9*is a target of FOXP2 in either developing or adult human brain (Gene specific Chipsq (Spiteri et al 2007, Vernes et al 2007; Text-based mining (Han et al 2015)). Alternatively, an indirect interaction could explain the observation, if FOXP2 represses expression of a key palmitoylated target. One candidate target for both genes could be *CNTNAP2*, another gene associated with language deficits, which is predicted to be palmitoylated based on protein sequence (SwissPalm database). Connectome analysis of cases with *F0XP2* mutations could provide further lines of evidence regarding similarity in neurodevelopmental pathology.

### Comparison to structural connectome studies in other disorders

The influence of *ZDHHC9* mutation on structural brain organisation shows both similarities and differences to published results on groups with a similar phenotype or genetic mechanisms.

Several studies investigated structural network organisation in Fragile-X syndrome. Similar to *ZDHHC9* mutation (Raymond et al., 2007), Fragile-X syndrome is a cause of X-linked intellectual disability. Leow et al. investigated local and global properties of the white matter connectome in Fragile-X syndrome (FXS). FXS is caused by CGG trinucleotide repeats in the Fragile-X mental retardiation 1 (*FMR1*) gene on the X chromosome (Belmonte & Bourgeron, 2006). Leow and colleagues reported an association between the number of trinucleotide repeats in the *FMR1* gene and global network efficiency in male premutation carriers (Leow et al., 2014) as well as local differences in efficiency and clustering coefficient in left temporal nodes (also see Bruno et al. 2016). Our results for *ZDHHC9* also indicated a reduction in global efficiency of the structural network similar to that reported for FXS, suggesting that this observation relates non-specifically to low IQ. However, topographical analysis of clustering coefficient and local efficiency indicated reductions in the parietal and frontal lobe in the *ZDHHC9* group, whereas reductions in temporal areas were less pronounced or were statistically indistinguishable from the control group. In other words, mutations in *ZDHHC9* and *FXS* show a convergent reduction in global network efficiency, but different local patterns of efficiency and clustering coefficient that distinguish the groups.

Rolandic epilepsy is another relevant neurodevelopmental condition for comparison due to the overlapping phenotype of expressive language deficits and epilepsy with centro-temporal spikes that were also observed in the carriers of *ZDHHC9* mutation (Baker et al., 2015). A study by Besseling and colleagues identified a reduction in structural white matter connectivity of the Perisylvian system, including the left inferior frontal, supramarginal, and postcentral gyrus (Besseling et al., 2013a). Studies of functional connectivity indicated reduced integration of these areas and delayed convergence of structural and functional connectivity in RE. (Besseling et al. 2013b & 2014). Further, graph theoretical analysis of the functional connectome indicated reduced clustering coefficient and local efficiency in areas of the parietal and frontal lobe in RE similar to the findings of structural connectivity differences in the current study (Xiao et al., 2015). In summary, studies of functional and structural connectivity in a neurodevelopmental condition of mixed aetiology with a similar phenotype to *ZDHHC9* mutation showed reduced connectivity in areas of the parietal and frontal lobe akin to the structural connectivity changes observed in the current investigation. We are not aware of another connectome analysis of a developmental language disorder (either of known or unknown origin) against which to compare the results of our study.

## Limitations

The findings of the current investigation are associated with some limitations. First, the sample size of the current study was inherently limited by the rarity of single-gene mutations associated with XLID. Further, group comparisons in the current study were made relatively to an age-matched typical control group rather than an IQ-matched control. Comparison to a mixed-aetiology low IQ group would be confounded by the association between intellectual disability and diverse structural abnormalities, whilst comparison to an alternative single-aetiology low IQ group with its own profile of neuroanatomical abnormality could be equally difficult to interpret.

In addition to limitations associated with the study sample, same caveats regarding the analysis methods need to be considered. A multitude of methods for structural connectome analysis of diffusion-weighted MR data have been reported in the literature (Qi 2015). There is currently no consensus on best practices or published rigorous comparisons across different methods. Biological validation of methods would require extensive in vivo studies that are difficult to carry out. In the absence of biological validation, Zalesky (2010) and Qi (2015) systematically compared the influence of various choices in the processing pipeline on graph theoretical comparisons. Following the recommendations arising from these comparisons, the same scanning protocol, diffusion model (constrained spherical deconvolution), and parcellation scheme was used for all participants in the current study so that any group differences are unlikely to be explained by these factors. Additionally, ROI size may influence connectivity results, and is likely to differ between case and control groups. Several methods for correcting this effects have been used (Daducci 2012, Hagmann 2012). In the current study, streamlines were transformed to a common space to alleviate differences in ROI size while maintaining a standard definition of anatomical cortex parcellation (Horn 2016). Another choice concerns the measure of connection strength. The current investigation used FA as the measure of connection strength as this measure is a common measure used in diffusion-weighted imaging that is easy to interpret in contrast to alternative approaches that use streamline-based measures.

## Conclusion

Mutations in the *ZDHHC9* gene were associated with altered network properties, including reduction in mean clustering coefficient and global efficiency. Topological analysis indicated that reductions in the *ZDHHC9* group were most pronounced for frontal and temporo-parietal nodes. Further, comparison of graph theory metrics with *ZDHHC9* expression data obtained from the Allen Brain Human Brain repository indicated that nodal variation in the clustering coefficient and local efficiency related to higher expression of *ZDHHC9*. This relationship between gene expression and regional variation in white matter organisation may provide a mechanistic link between this particular gene and the disproportionate impact of this mutation of speech and language development. In short, white matter networks may represent an important intermediate phenotype to understand the effect of genetic mutations on cognitive development.

